# Identification, Mapping and Relative Quantitation of SARS-CoV-2 Spike Glycopeptides by Mass-Retention Time Fingerprinting

**DOI:** 10.1101/2020.07.24.217562

**Authors:** R. Chalk, W. Greenland, T. Moreira, J. Coker, S.M.M Mukhopadhyay, E. Williams, C. Manning, T. Bohstedt, R. McCrorie, A. Fernandez-Cid, N.A. Burgess-Brown

**Affiliations:** Centre for Medicines Discovery, ORCRB, Oxford University, OX3 7DQ, UK; Agilent Technologies, Lakeside, Cheadle Royal Business Park, Cheadle, Cheshire, SK8 3GR, UK

**Keywords:** SARS-CoV-2, Spike, RBD, Glycoprotein, Glycopeptide, Glycan, Mass Spectrometry, HPLC, Database

## Abstract

We describe a novel analytical method for rapid and robust identification, mapping and relative quantitation of glycopeptides from SARS-CoV-2 Spike protein. The method may be executed using any LC-TOF mass spectrometer, requires no specialised knowledge of glycan analysis and makes use of the differential resolving power of reversed phase HPLC. While this separation technique resolves peptides with high efficiency, glycans are resolved poorly, if at all. Consequently, glycopeptides consisting of the same peptide bearing different glycan structures will all possess very similar retention times and co-elute. While this has previously been viewed as a disadvantage, we show that shared retention time can be used to map multiple glycan species to the same peptide and location. In combination with MSMS and pseudo MS3, we have constructed a detailed mass-retention time database for Spike. This database allows any ESI-TOF equipped lab to reliably identify and quantify spike glycans from a single overnight elastase protein digest in less than 90 minutes.

## Introduction

Glycosylation is known to play an important role in the efficacy and antigenicity of therapeutic proteins [1–3]. The current SARS-CoV-2 pandemic has spurred urgent research, much of it devoted to preparing vaccines, therapeutic antibodies or antibody tests based on Spike protein, the virus’s primary surface antigen [4]. This 145 kDa protein forms a trimer [5] with each subunit bearing twenty-two potential N-linked glycosylation sites and two O-linked sites of which approximately seventeen are occupied [5]. The unusually heavy and complex glycosylation observed in Spike protein is believed to play an important role in the pathogenicity of SARS-CoV-2 by mimicking host cell glycans and allowing the virus to evade the normal immune response [6]. Analysis of expressed Spike protein by mass spectrometry presents unique challenges in terms of its size and the number and complexity of its glycans. These challenges have been commendably met to date by laboratories with wide experience in glycan analysis and access to very sensitive, high-end nano-LC-MSMS mass spectrometers [1, 7–9]. However, in our laboratory and in others a rapid and more robust methodology is needed for routine analysis of different batches of expressed Spike protein. In addition, any method which is reliant on LC-MSMS of glycopeptides may not necessarily detect specific glycans which fail to fragment under the conditions selected. LC-MS, by contrast, generates a mass, retention time and relative abundance for all ionizable species. We have developed a simple Mass-Retention Time Fingerprinting (MRTF) method for rapid and robust identification, mapping and relative quantitation of Spike glycans. Overnight digestion using a single enzyme followed by a 65-minute LC-MS run using any accurate mass instrument are the only experimental requirements. The resulting LC-MS data contains accurate mass, retention time and relative abundance values for each glycopeptide component. This dataset needs only to be matched against the pre-existing Spike glycopeptide database reported here, as shown in Figure 1. We describe this method as “analytical mode”, which is both conceptually simple to understand, and straightforward to implement in a typical mass spectrometry laboratory. For scientific completeness, we also describe the “discovery mode” which we have used to generate the data for our Mass-Retention Time Fingerprinting database. We stress, however, that there is no requirement for users to duplicate this discovery mode. The analytical mode in conjunction with the database we have provided is all that is necessary to characterise Spike glycans by MRTF.

**Figure 1.**
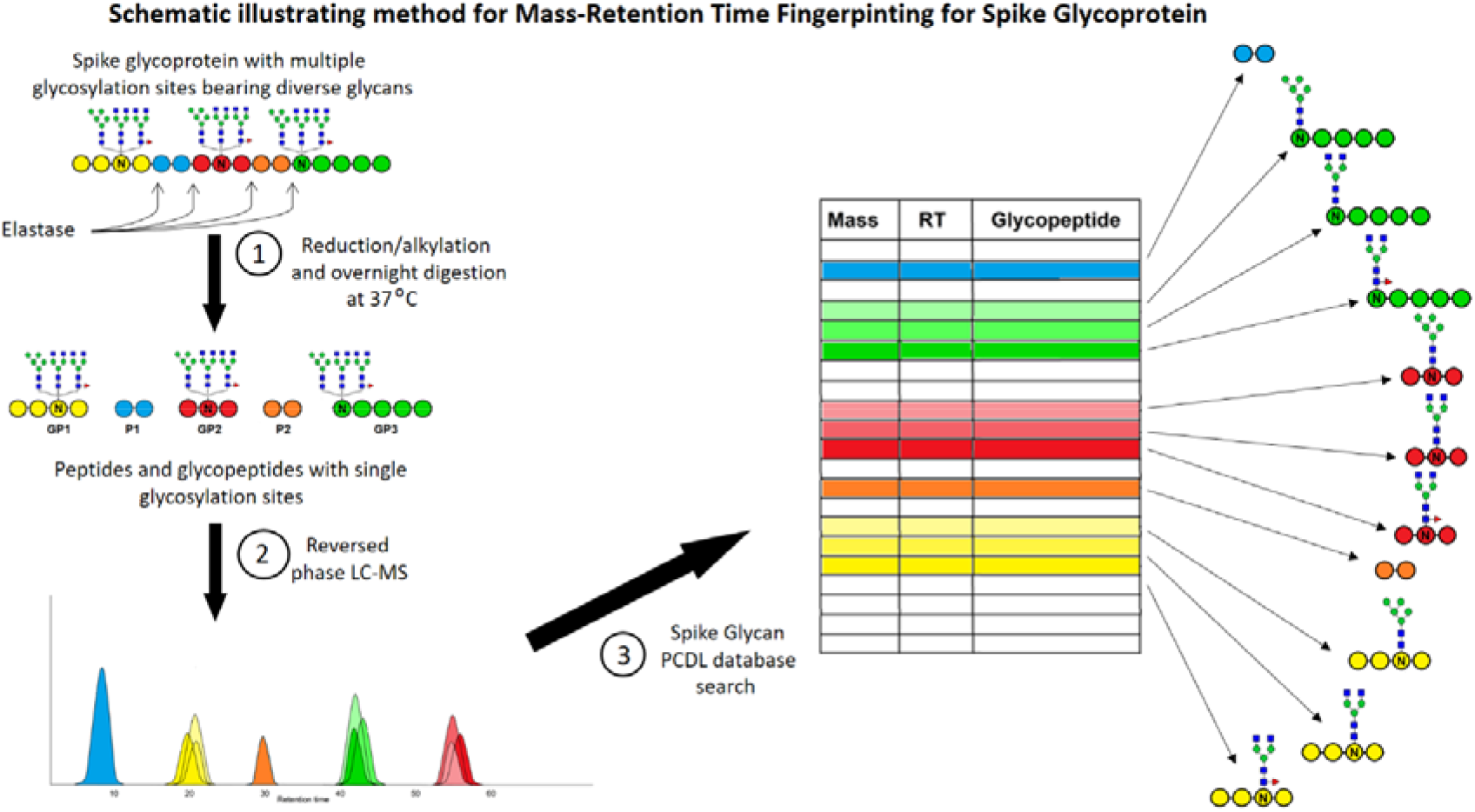
Mass-Retention Time Fingerprint analysis of Spike glycans

## Results

Combined extracted ion chromatograms are illustrated in Figure 2. It can be seen that some species (peptides) are completely resolved, while many other species (glycans) co-elute.

**Figure 2.**
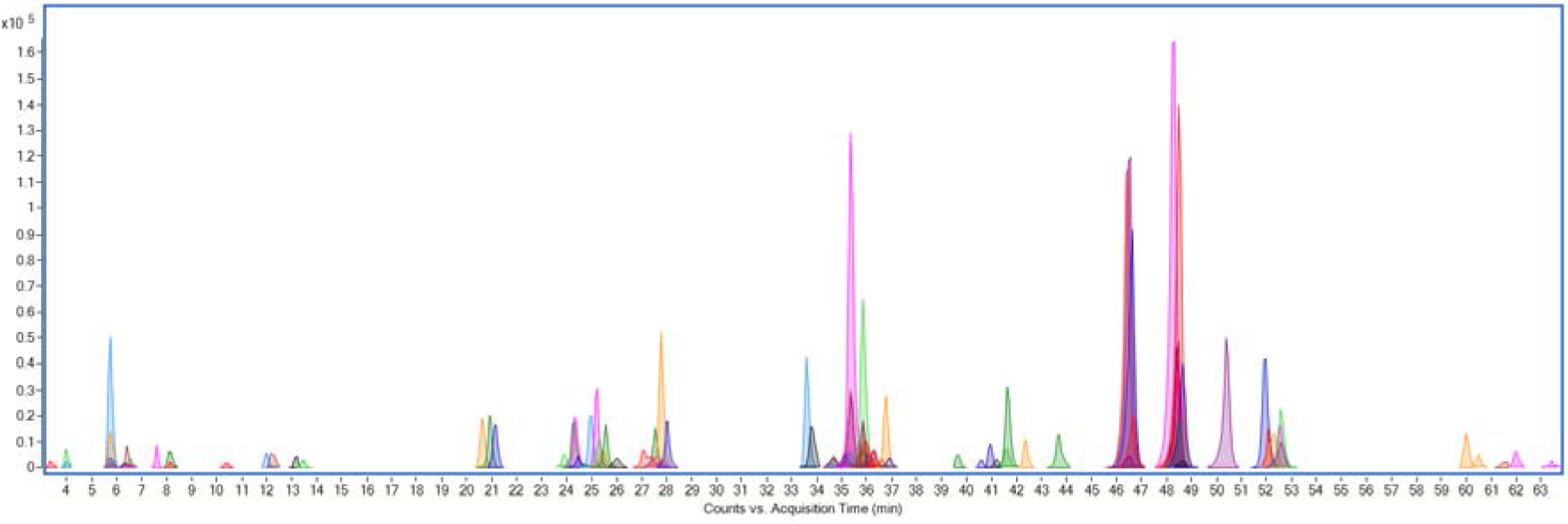
Combined Extracted Ion Chromatogram (EIC) for 140 observed Spike glycopeptides separated by reversed phase LC-MS and identified by Mass-Retention Time Fingerprinting (MRTF)

**Figure 3.**
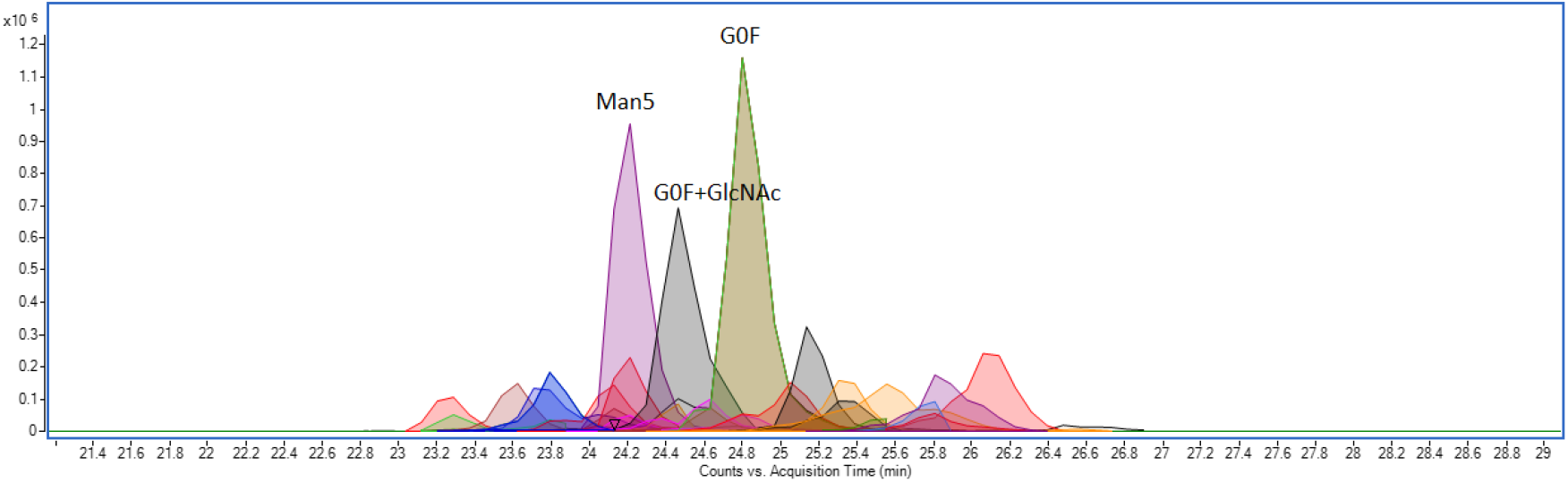
Combined Extracted Ion Chromatogram (EIC) for 27 isoforms of glycopeptide GEVFNAT (N343) within +/− 2 min retention time window from RBD. Only three glycans are labelled, the remainder are listed in the accompanying table 2, below.

We observed one hundred and forty glycopeptides by LC-MS. These are recorded grouped by ascending retention time in Tables 1a and 1b along with accurate masses, peptide sequences, glycan assignments and a key to the glycan structures. The location of each glycopeptide series on Spike is indicated in the first column. It may be seen that all observed glycans for the same peptide occur within a four minute retention time window. Accurate mass and estimated retention time are included for a further three hundred and six glycopeptides.

**Table 1a.**
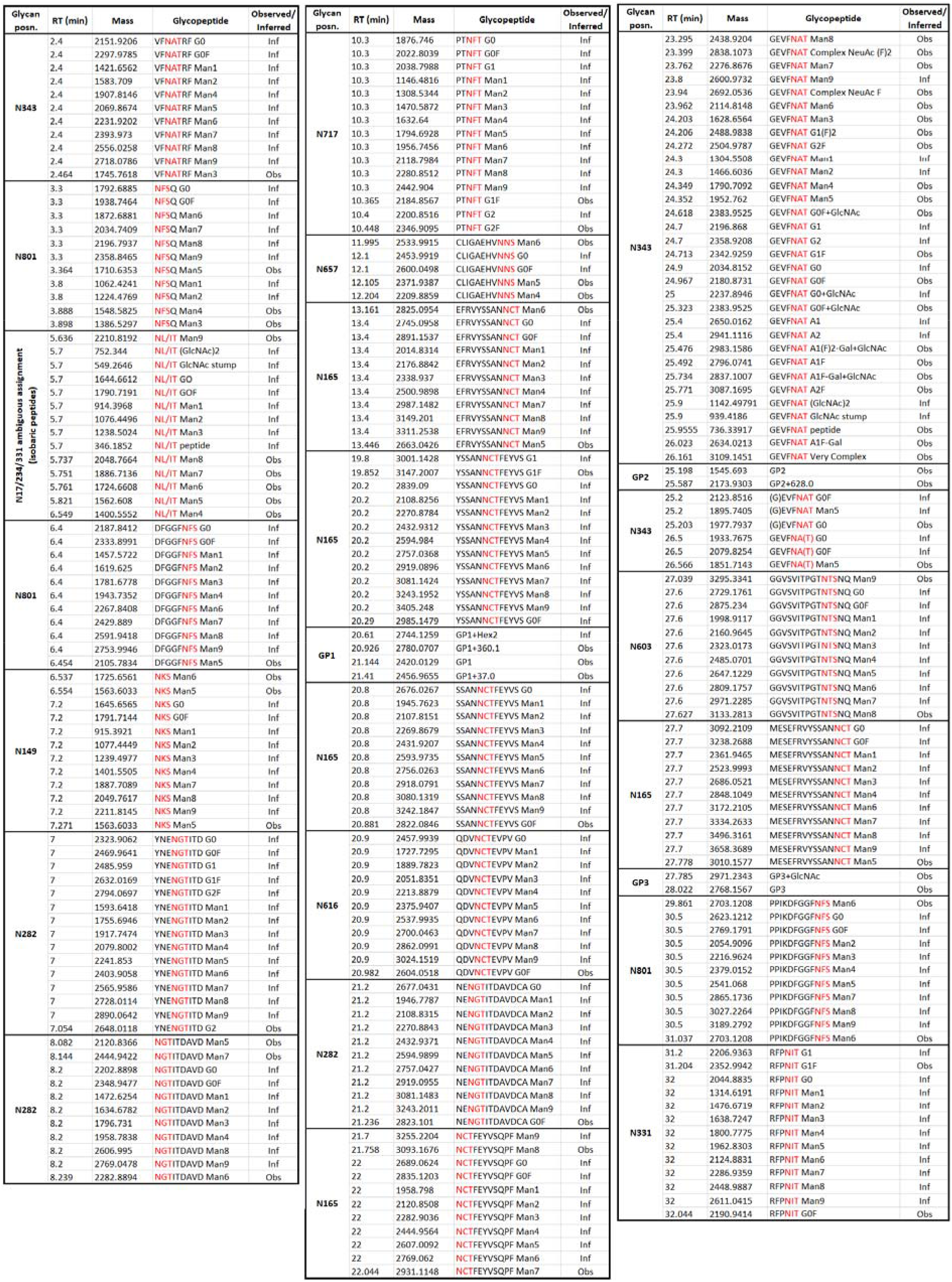
Spike elastase glycopeptide mass retention time database (PCDL) containing data for 140 observed glycopeptides and data for a further 306 inferred glycopeptides (RT 2-32 min)

**Table 1b.**
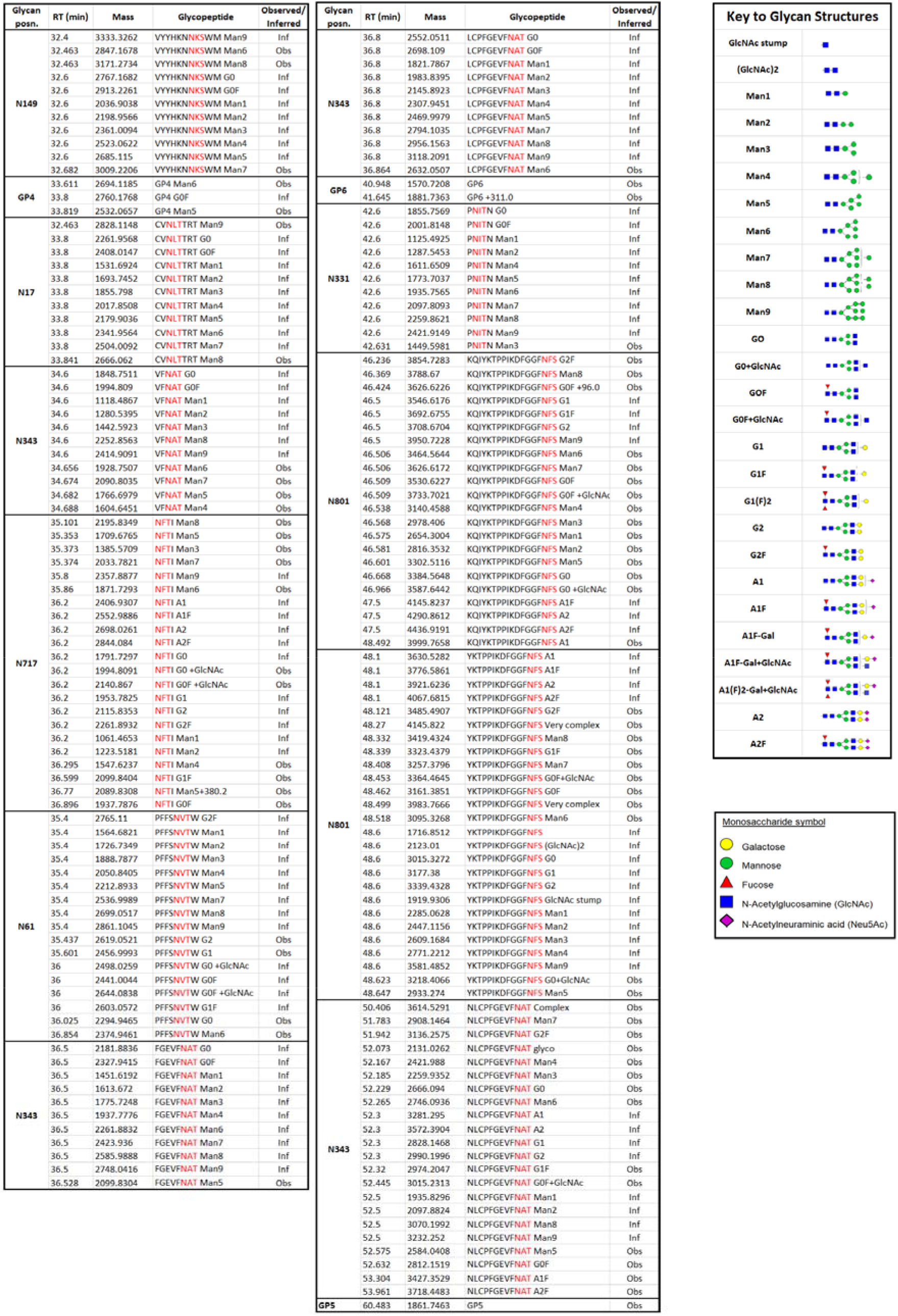
Spike elastase glycopeptide mass retention time database (PCDL) containing data for 140 observed glycopeptides and data for a further 306 inferred glycopeptides (RT 32-60 min and key)

**Table 2.** GEVFNAT glycopeptide (N343) isoforms from RBD shown in Figure 3

| Name | Mass | RT | Volume | ppm error | Name | Mass | RT | Volume | ppm error |
| --- | --- | --- | --- | --- | --- | --- | --- | --- | --- |
| GEVFNAT Complex NeuAc (F)2 | 2838.1078 | 23.26 | 733674 | -0.2 | GEVFNAT G1F | 2342.9180 | 24.62 | 1090359 | 3.4 |
| GEVFNAT Man8 | 2438.9113 | 23.30 | 744169 | 3.7 | GEVFNAT G0 | 2034.8089 | 24.80 | 950868 | 3.1 |
| GEVFNAT Man7 | 2276.8615 | 23.61 | 1867525 | 2.7 | GEVFNAT G0F | 2180.8688 | 24.82 | 12980259 | 2.0 |
| GEVFNAT Complex NeuAc F | 2692.0523 | 23.77 | 1266219 | 0.5 | GEVFNAT A1(F)2-Gal+GlcNAc | 2983.1427 | 24.91 | 732781 | 5.3 |
| GEVFNAT Man6 | 2114.8091 | 23.80 | 2432950 | 2.7 | (G)EVFNAT G0 | 1977.7887 | 25.05 | 2947447 | 2.5 |
| GEVFNAT G1(F)2 | 2488.9768 | 24.09 | 1292274 | 2.8 | GEVFNAT G0F+GlcNAc | 2383.9464 | 25.16 | 3836871 | 2.6 |
| GEVFNAT G2F | 2504.9755 | 24.14 | 1007345 | 1.3 | GEVFNAT A1F | 2796.0641 | 25.33 | 1512501 | 3.6 |
| GEVFNAT Man4 | 1790.7051 | 24.20 | 3040952 | 2.3 | GEVFNAT A1(F)2-Gal+GlcNAc | 2983.1444 | 25.34 | 1320994 | 4.8 |
| GEVFNAT Man5 | 1952.7583 | 24.20 | 12492713 | 1.9 | GEVFNAT A1F-Gal | 2634.0105 | 25.51 | 608358 | 4.1 |
| GEVFNAT Man3 | 1628.6504 | 24.20 | 666739 | 3.7 | GEVFNAT A2F | 3087.1629 | 25.77 | 805142 | 2.2 |
| GEVFNAT G1F | 2342.9183 | 24.37 | 1333546 | 3.3 | GEVFNAT A1F | 2796.0644 | 25.80 | 541989 | 3.5 |
| GEVFNAT G0F | 2180.8638 | 24.45 | 953034 | 4.3 | GEVFNAT A1F-Gal | 2634.0125 | 25.86 | 3692097 | 3.4 |
| GEVFNAT G0F+GlcNAc | 2383.9436 | 24.47 | 9038631 | 3.7 |  |  |  |  |  |

The complete Spike PCDL database is available to download in .cdb or .xlsx format here: https://zenodo.org/record/3958218#.Xxn_BChKhoY

Figure 4 illustrates a complete glycan fragmentation series for RBD glycopeptide GEVFNAT-Man5 showing the peptide stump (GEVFNAT-GlcNAc) and mannose ladders. Calculated mass errors are shown in table 3.

**Figure 4.**
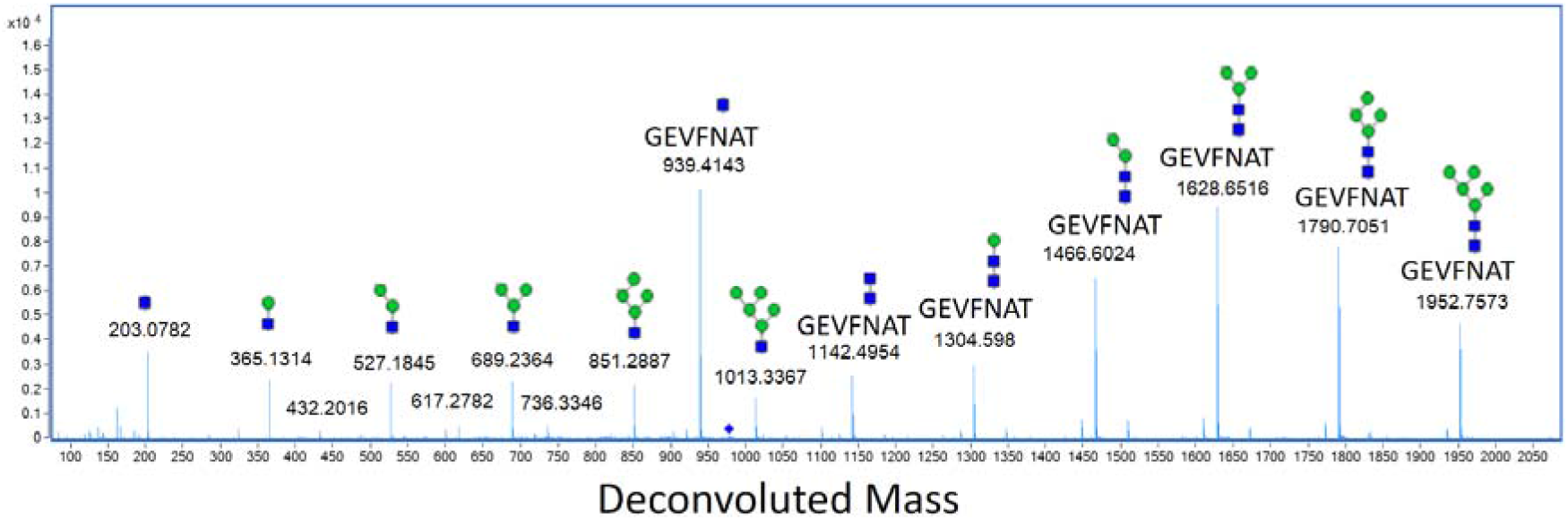
Complete glycan fragmentation series for RBD glycopeptide GEVFNAT-Man5 (N343)

**Table 3.** Glycan assignment and mass errors (parts per million) for RDB glycopeptide GEVFNAT-Man5

| Deconvoluted Mass Obs | Formula | Mass Calc | ppm | Assignment |
| --- | --- | --- | --- | --- |
| 203.0782 | C<sub>8</sub>H<sub>13</sub>N O<sub>5</sub> | 203.0794 | -5.9 | GlcNAc |
| 365.1314 | C<sub>8</sub>H<sub>13</sub>N O<sub>5</sub> (C<sub>6</sub>H<sub>10</sub>O<sub>5</sub>)<sub>1</sub> | 365.1322 | -2.2 | GlcNAc(Man)<sub>1</sub> |
| 527.1845 | C<sub>8</sub>H<sub>13</sub>N O<sub>5</sub> (C<sub>6</sub>H<sub>10</sub>O<sub>5</sub>)<sub>2</sub> | 527.1850 | -0.9 | GlcNAc(Man)<sub>2</sub> |
| 689.2364 | C<sub>8</sub>H<sub>13</sub>N O<sub>5</sub> (C<sub>6</sub>H<sub>10</sub>O<sub>5</sub>)<sub>3</sub> | 689.2364 | 0.0 | GlcNAc(Man)<sub>3</sub> |
| 851.2887 | C<sub>8</sub>H<sub>13</sub>N O<sub>5</sub> (C<sub>6</sub>H<sub>10</sub>O<sub>5</sub>)<sub>4</sub> | 851.2907 | -2.3 | GlcNAc(Man)<sub>4</sub> |
| 1013.3367 | C<sub>8</sub>H<sub>13</sub>N O<sub>5</sub> (C<sub>6</sub>H<sub>10</sub>O<sub>5</sub>)<sub>5</sub> | 1013.3435 | -6.7 | GlcNAc(Man)<sub>5</sub> |
| 939.4143 | C<sub>32</sub>H<sub>48</sub>N<sub>8</sub>O<sub>12</sub> (C<sub>8</sub>H<sub>13</sub>N O<sub>5</sub>) | 939.4185 | -4.5 | GEVFNAT (GlcNAc) |
| 1142.4954 | C<sub>32</sub>H<sub>48</sub>N<sub>8</sub>O<sub>12</sub> (C<sub>8</sub>H<sub>13</sub>N O<sub>5</sub>)<sub>2</sub> | 1142.4979 | -2.2 | GEVFNAT (GlcNAc)<sub>2</sub> |
| 1304.5498 | C<sub>32</sub>H<sub>48</sub>N<sub>8</sub>O<sub>12</sub> (C<sub>8</sub>H<sub>13</sub>N O<sub>5</sub>)<sub>2</sub> (C<sub>6</sub>H<sub>10</sub>O<sub>5</sub>)<sub>1</sub> | 1304.5507 | -0.7 | GEVFNAT (GlcNAc)<sub>2</sub> (Man)<sub>1</sub> |
| 1466.6024 | C<sub>32</sub>H<sub>48</sub>N<sub>8</sub>O<sub>12</sub> (C<sub>8</sub>H<sub>13</sub>N O<sub>5</sub>)<sub>2</sub> (C<sub>6</sub>H<sub>10</sub>O<sub>5</sub>)<sub>2</sub> | 1466.6036 | -0.8 | GEVFNAT (GlcNAc)<sub>2</sub> (Man)<sub>2</sub> |
| 1628.6516 | C<sub>32</sub>H<sub>48</sub>N<sub>8</sub>O<sub>12</sub> (C<sub>8</sub>H<sub>13</sub>N O<sub>5</sub>)<sub>2</sub> (C<sub>6</sub>H<sub>10</sub>O<sub>5</sub>)<sub>3</sub> | 1628.6564 | -2.9 | GEVFNAT (GlcNAc)<sub>2</sub> (Man)<sub>3</sub> |
| 1790.7051 | C<sub>32</sub>H<sub>48</sub>N<sub>8</sub>O<sub>12</sub> (C<sub>8</sub>H<sub>13</sub>N O<sub>5</sub>)<sub>2</sub> (C<sub>6</sub>H<sub>10</sub>O<sub>5</sub>)<sub>4</sub> | 1790.7092 | -2.3 | GEVFNAT (GlcNAc)<sub>2</sub> (Man)<sub>4</sub> |
| 1952.7573 | C<sub>32</sub>H<sub>48</sub>N<sub>8</sub>O<sub>12</sub> (C<sub>8</sub>H<sub>13</sub>N O<sub>5</sub>)<sub>2</sub> (C<sub>6</sub>H<sub>10</sub>O<sub>5</sub>)<sub>5</sub> | 1952.7620 | -2.4 | GEVFNAT (GlcNAc)<sub>2</sub> (Man)<sub>5</sub> |

In the pseudo MS3 experiment glycans were lost by in-source decay. GEVFNAT-GlcNAc was isolated in the quadrupole and fragmented in the collision cell. Sequence confirmation for the peptide stump GEVFNAT-GlcNAc is shown in Figure 5 with mass errors calculated in Table 4.

**Figure 5.**
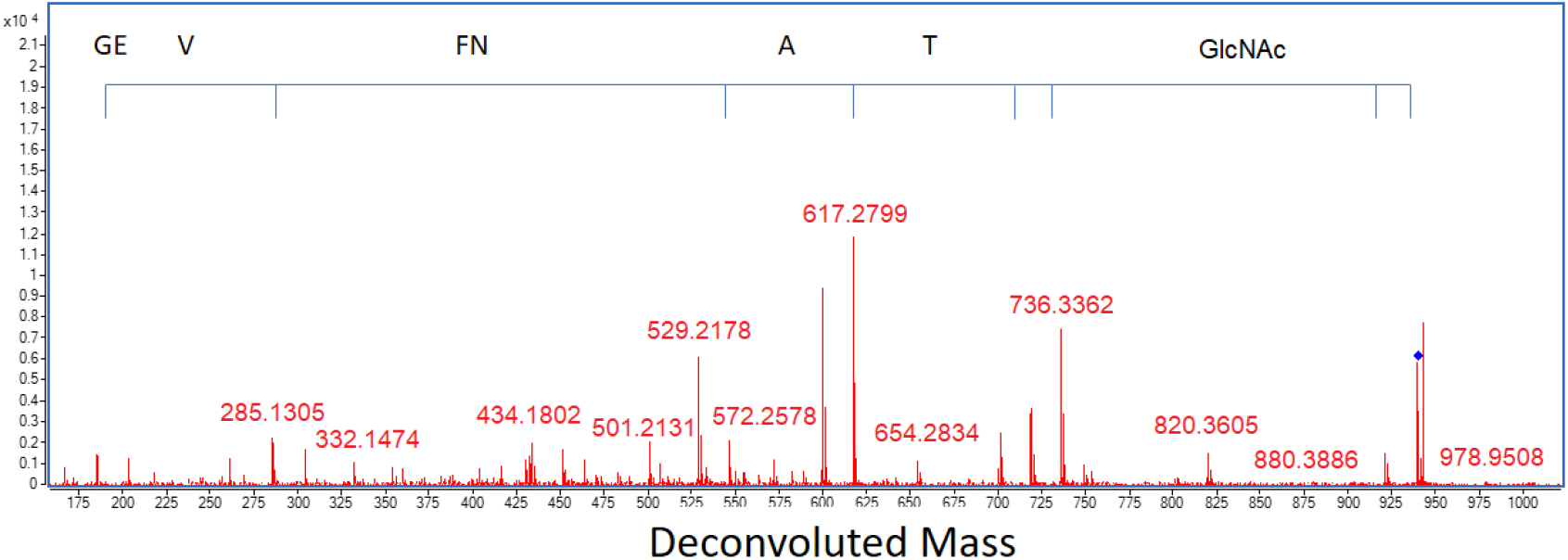
Pseudo MS3 fragmentation analysis of RBD glycopeptide stump GEVFNAT-GlcNAc (N343)

**Table 4.** Pseudo MS3 fragment ion assignment and mass errors (parts per million) for RBD glycopeptide stump GEVFNAT-GlcNAc (N343)

| Deconvoluted Observed Mass | Formula | Calculated Mass | Error (ppm) | Fragment ion assignment | Peptide Sequence |
| --- | --- | --- | --- | --- | --- |
| 186.0645 | C<sub>7</sub>H<sub>10</sub>N<sub>2</sub>O<sub>4</sub> | 186.0641 | 2.1 | b2 | GE |
| 285.1305 | C<sub>12</sub>H<sub>19</sub>N<sub>3</sub>O<sub>5</sub> | 285.1325 | -7.0 | b3 | GEV |
| 546.2410 | C<sub>25</sub>H<sub>34</sub>N<sub>6</sub>O<sub>8</sub> | 546.2438 | -5.1 | b5 | GEVFN |
| 617.2799 | C<sub>28</sub>H<sub>39</sub>N<sub>7</sub>O<sub>9</sub> | 617.2809 | -1.6 | b6 | GEVFNA |
| 718.3257 | C<sub>32</sub>H<sub>46</sub>N<sub>8</sub>O<sub>11</sub> | 718.3286 | -4.0 | b7 | GEVFNAT |
| 736.3362 | C<sub>32</sub>H<sub>48</sub>N<sub>8</sub>O<sub>12</sub> | 736.3392 | -4.1 | y7 | GEVFNAT |
| 939.4174 | C<sub>32</sub>H<sub>48</sub>N<sub>8</sub>O<sub>12</sub> C<sub>8</sub>H<sub>13</sub>N O<sub>5</sub> | 939.4185 | -1.2 | M GlcNAc | GEVFNAT GlcNAc |
| 921.4064 | C<sub>32</sub>H<sub>46</sub>N<sub>8</sub>O<sub>11</sub> C<sub>8</sub>H<sub>13</sub>N O<sub>5</sub> | 921.408 | -1.7 | M GlcNAc - H<sub>2</sub>O | GEVFNAT GlcNAc |
| 820.3605 | C<sub>28</sub>H<sub>39</sub>N<sub>7</sub>O<sub>9</sub> C<sub>8</sub>H<sub>13</sub>N O<sub>5</sub> | 820.3603 | 0.2 | b6 GlcNAc - H<sub>2</sub>O | GEVFNA GlcNAc |
| 749.3213 | C<sub>25</sub>H<sub>34</sub>N<sub>6</sub>O<sub>8</sub> C<sub>8</sub>H<sub>13</sub>N O<sub>5</sub> | 749.3232 | -2.5 | b5 GlcNAc - H<sub>2</sub>O | GEVFN GlcNAc |
| 701.3021 | C<sub>32</sub>H<sub>43</sub>N<sub>7</sub>O<sub>11</sub> | 701.3021 | 0.0 | b7 - NH<sub>3</sub> | GEVFNAT |
| 600.2529 | C<sub>28</sub>H<sub>36</sub>N<sub>6</sub>O<sub>9</sub> | 600.2544 | -2.5 | b6 - NH<sub>3</sub> | GEVFNA |
| 529.2178 | C<sub>25</sub>H<sub>31</sub>N<sub>5</sub>O<sub>8</sub> | 529.2173 | 0.9 | b5 - NH<sub>3</sub> | GEVFN |

Intact mass measurement of fully glycosylated Spike was unsuccessful due to the polydispersity of its innumerable glycoforms and the resulting dilution of ion signal. However, the smaller receptor binding domain, bearing only two glycosylation sites did prove amenable to intact mass analysis. Figure 6 shows twenty-one glycoforms for intact RBD, of which ten major glycoforms could be assigned. This showed that the principal glycan species were Man5, G0F and G0F+GlcNAc which was in agreement with the glycopeptide analysis.

**Figure 6.**
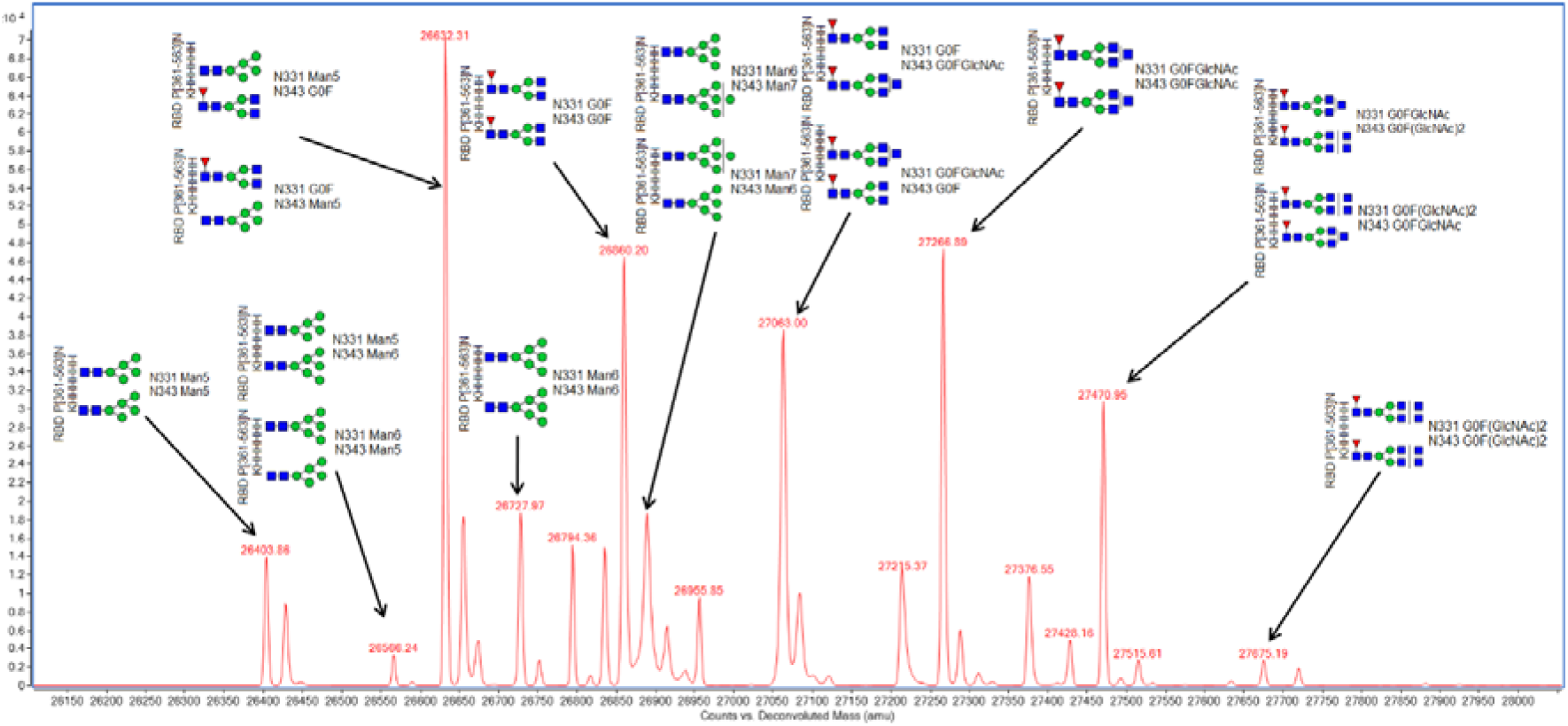
Intact mass analysis of RBD showing the principal glycan species Man5, G0F and G0F+GlcNAc in agreement with glycopeptide analysis. (This method cannot differentiate individual glycosylation sites, hence when two structures are possible, both are shown)

Elastase was chosen as a single digestion enzyme because it was judged to give the best chance of generating glycopeptides with a single NXS/T motif, essential for unambiguous glycan mapping. For non-glycosylated Spike peptides, elastase generated 63 high quality MSMS hits and 26% coverage allowing for five missed cleavages. The same data searched for non-specific cleavage gave 135 high quality MSMS hits and 48% coverage allowing for twenty missed cleavages. Elastase itself contains 2 NXS/T motifs. We therefore prepared elastase only, at x10 the usual concentration, searched the resulting LC-MS data using the PCDL as a control, and no hits were found. The Spike protein LC-MS data did contain a small number of elastase autodigestion peptides.

## Methods

### Cloning, expression and purification of Spike

The gene encoding amino acids 1-1208 of the SARS-CoV-2 Spike glycoprotein ectodomain (S), with mutations of RRAR > GSAS at residues 682-685 (to remove the furin cleavage site) and KV > PP at residues 986-987 (to stabilise the protein), was synthesised with a C-terminal T4 fibritin trimerization domain, HRV 3C cleavage site, 8xHis tag, and Twin-Strep-tag [5]. The construct was sub-cloned into pHL-sec [10] using the AgeI and XhoI restriction sites and the sequence was confirmed by sequencing. Recombinant Spike was produced in *Expi293F*^*TM*^cells by transient transfection with purified DNA (0.5 mg/L cells) using a 1:6 DNA:L-PEI ratio, mixed in minimal medium, and sodium butyrate as an additive. Cells were grown in suspension in *FreeStyle293^TM^* medium with shaking at 150 rpm in 2 L smooth roller bottles, filled with 0.5 L cells at 2 e^6^/mL per bottle at 30°C with 8% CO_2_ and 75% humidity. Supernatants from transfected cells were harvested 3-days post-transfection by centrifugation. Clarified supernatant was mixed with Ni^2+^ IMAC *Sepharose*^®^ *6 Fast Flow* (*GE*; 2 mL bed volume per L of supernatant) at room temperature for 2 h. Using a gravity flow column, resin was collected and washed stringently with 50 CV each of base buffer (1X PBS), WB25 (BB + 25 mM imidazole), and WB40 (BB+ 40 mM imidazole), followed by elution with EB (0.30 M imidazole in 1X PBS). Protein was dialyzed into 1X PBS using *SnakeSkin*^*TM*^ 3,500 MWCO dialysis tubing, concentrated to 1 mg/mL using a 100,000 MWCO *VivaSpin* centrifugal concentrator (*GE*), and centrifuged at 21,000 x *g* for 30 min to remove aggregates. The trimeric Spike protein was flash frozen in LN_2_ and stored at −80°C until use. Final purified yield was 1 mg of Spike protein per L of transfected cells.

### Cloning, expression and purification of Receptor Binding Domain

The receptor binding domain (RBD; aa 330-532) of SARS-CoV-2 Spike (Genbank MN908947) was inserted into the pOPINTTGneo expression vector fused to an N-terminal signal peptide and a C-terminal 6xHis tag [11]. RBD was produced by transient transfection in *Expi293F*^*TM*^ cells (*ThermoFisher Scientific*, UK) using purified DNA (1.0 mg/L cells), a 1:3 DNA:L-PEI ratio, and sodium butyrate as an additive. Cells were grown in suspension in *FreeStyle293*^*TM*^ expression medium at 37°C with 8% CO_2_ and 75% humidity. Supernatants from transfected cells were harvested 3-days post-transfection and the supernatant was collected by centrifugation. Clarified supernatant was incubated with 5 mL of Ni^2+^ IMAC *Sepharose*^®^ *6 Fast Flow* (*GE*) at room temperature for 2 h. Using gravity flow, resin was washed with 50 CV of base buffer (1X PBS) and 50 CV of WB (1X PBS + 25 mM imidazole) before elution with EB (0.5 M imidazole in 1X PBS). Protein was concentrated using a 10,000 MWCO *Amicon Ultra-15* before application to a *Superdex 75* 16/600 column pre-equilibrated with 1X PBS pH 7.4. Peak monomeric fractions were pooled and concentrated to 2 mg/mL, flash frozen in LN_2_, and stored at −80°C until use. Final purified yield was >15 mg RBD per L of transfected cells.

### Sample preparation

SARS-CoV-2 Spike or RBD-6H at 1 mg/mL in PBS were prepared in aliquots of either 20 μL or 80 μL and diluted 1 in 3 in 100 mM ammonium bicarbonate, pH 8.0, followed by reduction by addition of 1, 4 Dithiothreitol (DTT) to 5 mM and incubation 37°C for 1 h. Next, the protein was alkylated by addition of iodoacetamide (IAA) to 15 mM and incubation in the dark for 30 min. This was followed by overnight digestion using elastase (*Promega*) at a ratio of 1:20 (w/w). The following day, the supernatant was dried using a rotary evaporator, and re-suspended in 60 μL of 0.1% formic acid for injection into the LC-MS.

### ‘Analytical mode’ LC-MS glycopeptide data acquisition

LC-MS ‘analytical mode’ was performed using a *1290 Infinity* UHPLC coupled to a *G6530A* ESI QTOF mass spectrometer (*Agilent Technologies*). TOF and quadrupole were calibrated prior to analysis and the reference ion 922.0098*m/z* was used for continuous mass correction. Sample was introduced using a 50 μL full-loop injection. Reversed phase chromatographic separation was achieved using an *AdvancedBio Peptide* reversed phase 2.7 μm particle, 2.1 mm × 100 mm column 655750-902 (*Agilent Technologies*). Mobile phase A was 0.1% formic acid in water and mobile phase B 0.1% formic acid in methanol (*Optima* LC-MS grade, *Fisher*). Initial conditions were 5% B and 0.200 mL/min flow rate. A linear gradient from 5% B - 60% B was applied over 60 min, followed by isocratic elution at 100% B for 2 min returning to initial conditions for a further 2 min. Post time was 10 min. MS source parameters were drying gas temperature 350°C, drying gas 8 L/min, nebulizer 30 psi, capillary 4000 V, fragmentor 150 V. MS spectrum range was 100 – 3200 *m/z* (centroid only), 2 GHz Extended Dynamic range, with the instrument in positive ion mode.

### LC-MSMS glycopeptide data acquisition ‘discovery mode’

LC-MSMS ‘discovery mode’ was performed as described above, with the following changes: Soft CID collision energy parameters for MSMS were slope 1.0, intercept 0 using argon as the collision gas (if using nitrogen slope 2.0, intercept 0) were used to favour glycan fragmentation over peptide fragmentation for glycopeptides. Sufficient non-glycosylated peptides were fragmented to give reasonable sequence coverage. Care was taken to reduce sodium and potassium contamination where possible and Tris buffers were avoided as these adducts interfere with glycopeptide analysis.

### LC-MS glycopeptide data analysis ‘analytical mode’

Analysis only required retention time and accurate mass data using the Spike PCDL database created as described below. This is possible either using the *Agilent* software described, software provided by other vendors, or by manual inspection. In our case, we used *Masshunter Qualitative Analysis* version B.07 (*Agilent Technologies*) and the Molecular Feature Extraction tool to extract H+, Na+ and K+ adducts and charge states +1 to +5. Briefly, this tool identifies and associates common spectral features such as carbon isotopes, adducts and multiple charge states as belonging to same Compound (peptide) by virtue of sharing the same accurate mass and retention time, then combines these features together to give a mass, retention time and volume for each compound. Compounds were then searched against Spike PCDL using a mass error window +/− 10 ppm and a retention time window +/− 2 min. Some filtering of the data was used to reduce the number of compounds and thence speed-up the PCDL search. Relative quantitation of each glycan on a particular glycopeptide could then be assessed.

### LC-MSMS glycopeptide analysis “discovery mode”

Construction of the Spike glycopeptide mass-retention time database (“discovery mode”) was more complex and time-consuming, but once constructed and made available to the scientific community, there is no further need to repeat this step. By using reverse phase HPLC, glycopeptides are separated by the relatively hydrophobic peptide moiety, whereas the associated hydrophilic glycans are grouped together by retention time as illustrated in figure 7.

**Figure 7.**
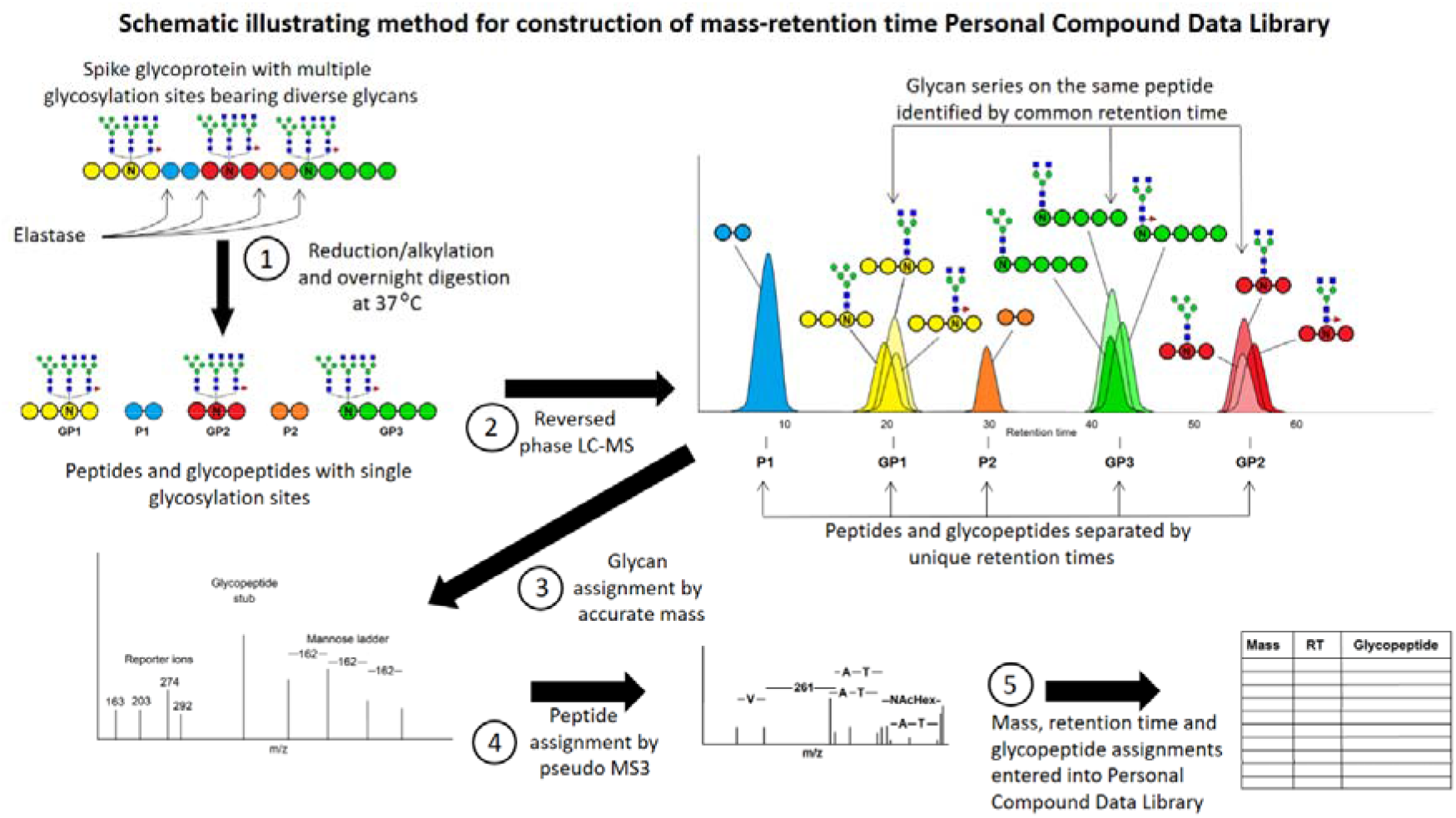
LC-MSMS “discovery” mode used to generate the Spike glycopeptide Mass-Retention Time PCDL database

Initial LC-MSMS discovery mode data for incorporation into a glycopeptide PCDL was performed using *Masshunter Qualitative Analysis* with *Bioconfirm* B.07.00 (*Agilent Technologies*). Compounds were identified using the Find by Molecular Feature (MFE) tool looking for H+, Na+ and K+ adducts and charge states +2 to +5. The results were filtered to remove compounds <1000 Da (too small to be glycopeptides). Compound MSMS spectra were screened manually for the following oxonium reporter ions: Hex *m/z* 163.0601, HexNAc *m/z* 204.0866, HexHexNAc *m/z* 366.1395, Neu5Ac *m/z* 274.0921/ *m/z* 291.0949 and/or a Hexose ladder _δ_M 162.0528 Da. High quality *m/z* spectra were deconvoluted to neutral mass spectra with glycan *de novo* interpretation performed manually. Once a glycopeptide had been identified, it was entered into a personal compound data library database (PCDL, *Agilent Technologies*) as a mass and retention time. In addition, the database made use of known mammalian N-linked glycan processing. After the initial glycopeptide identification, other processed glycopeptides, which were considered likely to also be present, were added to the database at the same retention time and with a calculated mass. For example, if a glycopeptide with Man5 was identified by MSMS, Man1-9 and G0/F were added at the same retention time. If these glycans were subsequently found in the data, their actual retention times were updated, and the next round of processing to more complex glycans was added, in order to produce the most comprehensive PCDL possible, while still being manageable. Processing order:

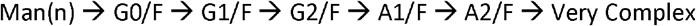

Valid glycan identifications resulted in a calculated peptide mass that could be matched to the sequence. Where high quality spectra were present, a peptide-GlcNAc stump was observed (Figure 4). This was used in a pseudo MS3 experiment with manual peptide *de novo* interpretation to confirm the peptide sequence (Figure 5). Mass data adjacent to the glycopeptide retention time was then searched for neutral differences corresponding to glycans, for example, Man5 → G0F or Man7 → G2F has a neutral delta mass of 228.1111 Da.

As expected, not all species could be matched to the sequence, presumably due to unexpected modifications. In this case, they were added to the database as ‘GP’ with an identifying number and as much information as could be extracted. Data for the most likely glycan was added to the PCDL, including a deconvoluted mass MSMS spectra were available, using nomenclature generating the most easily readable format.

A second round of glycopeptide discovery used *Bioconfirm* v10.0 data analysis software (*Agilent Technologies*). Sequences were matched by peptide accurate mass using the following parameters: peptide cleavage nonspecific, number of missed cleavages 20, N-linked modifications Man3, Man5-9, G0, G0F, G0F GlcNAc, G1, G1F, G2, G2F. Any peptide bearing the glycosylation motif NXS/T with two or more glycan hits within a retention time window +/− 2 min was added to the PCDL, excepting missed cysteine alkylations.

In-source fragmentation due to glycopeptide ions absorbing excess energy could be identified in the MS by searching extracted ion chromatograms (EICs) of the oxonium reporter ions and also by related glycopeptides appearing with exactly the same retention times. Both were observed infrequently and at manageable levels.

### Intact mass analysis

Concentrated protein samples were diluted to 0.02 mg/mL in 0.1% formic acid and 50 μL was injected on to a 2.1 mm x 12.5 mm *Zorbax* 5 μm *300SB-C3* guard column (*Agilent Technologies*) housed in a column oven set at 40°C. The solvent system used consisted of 0.1% formic acid (solvent A) and 0.1% formic acid in methanol (solvent B). Chromatography was performed as follows: Initial conditions were 90% A and 10% B and a flow rate of 1.0 mL/min. A linear gradient from 10% B to 80% B was applied over 35 seconds. Elution then proceeded isocratically at 95% B for 40 seconds followed by equilibration at initial conditions for a further 15 seconds. The mass spectrometer was configured with the standard ESI source and operated in positive ion mode. The ion source was operated with the capillary voltage at 4000 V, nebulizer pressure at 60 psig, drying gas at 350°C and drying gas flow rate at 12 L/min. The instrument ion optic voltages were as follows: fragmentor 250 V, skimmer 60 V and octopole RF 250 V.

## Discussion

Glycoprotein analysis is difficult. It is either performed in biopharmaceutical laboratories with proprietary expertise of glycan analysis on simple glycoproteins, such as immunoglobulins, or performed by a handful of academic labs with experience of glycan discovery from complex glycoproteins. Many protein researchers choose to ignore it, manipulating cell lines such that they cannot process beyond Man5, or to remove glycans entirely by mutation at the glycosylation motif or enzymatically [12]. While this approach has its merits, it has exposed a serious weakness in analytical capability when faced with a pathogen such as SARS-CoV-2 whose ability to evade the immune system is dependent upon heavy and complex glycosylation.

We have chosen an approach relying on elastase digestion to generate glycopeptides bearing a single glycan but with a sufficient number of amino acid residues to enable chromatographic separation by reversed-phase HPLC, as well as confident identification by accurate mass or *de novo* sequencing. Our choice of reversed phase HPLC has excellent discrimination for short elastase peptides, whereas glycans show little or no interaction with the column. Thus, species originating from a single glycosylation site with the same peptide sequence but several different glycans, eluted with the same retention time and could be discriminated by mass spectrometry. We used reversed phase HPLC and MSMS to characterise as many glycopeptides as possible. Although this required complex and time-consuming data analysis, it needed only be performed once, with the goal of building an accurate mass-retention time database for all observed Spike glycopeptides. Provided the same HPLC column and mobile phase conditions are used, retention times should not vary significantly. Thus, working in the analytical mode we describe, glycan structure and peptide sequence is assigned confidently, by accurate mass and retention time alone. LC-MS data need only to be searched against the mass-retention time database, and peak areas recorded, to generate a complete characterisation of Spike glycans.

We believe the MRTF method described here has advantages over other approaches to Spike glycan analysis. Previous studies relied upon very expensive equipment and software unavailable in most analytical laboratories. Working in ‘analytical’ mode, all that is necessary is to reproduce the chromatography, hence our method is a generic one, which can be run using any HPLC coupled to any accurate mass instrument and is not restricted to specific proprietary data analysis software. We used PCDL and *Masshunter*, but MRTF analysis can be performed on any vendor software or manually. Moreover, it demands no specialised expertise in glycobiology, and is thus accessible to many more researchers. Some published methods require multiple specific endoproteases, some of which cannot be readily sourced. Our method uses a single enzyme, elastase, which is inexpensive and widely available. Nor does it rely on glycosidases, which may not work efficiently and do not cleave O-linked glycans.

Our data contains an excess of glycopeptides with the motif (y)nNxS/T. This appears to be a very convenient function of elastase on glycopeptides, because the presence of the motif at the C-terminus facilitates *de novo* sequencing. We would be interested to know if this cleavage bias towards the C-terminus of the glycan motif is reproducible in other labs and whether it indicates steric hindrance within the elastase enzyme structure. If such bias is real, then these peptides are less likely to be a false positive result.

Receptor binding domain (RBD) from Spike protein is of interest in many labs for development of serological tests or neutralising antibodies. Because the yield of RBD was five times higher than Spike and more was initially available, we used it for method optimisation, and since it bears only two glycosylation sites which are also present on Spike, it functioned as a useful model. Consequently, N343 on glycopeptide GEVFNAT is over-represented in our demonstration PCDL. We consistently observed the same three major glycans (Man5, G0F and G0F+GluNAc) on this peptide and these were also in agreement with intact mass analysis of RBD protein as shown in Figure 6. On closer inspection, glycans up to A2F could also be observed at lower levels. We suspect that sufficiently detailed analysis may reveal all possible glycan structures with low abundance at all available sites. The most important would therefore be the top three to five glycans. If the complete complement of Spike protein glycoforms proves too challenging for a single analysis, this site, which is the most complete, would make a good proxy for total glycosylation.

We acknowledge that the mass-retention time fingerprinting method described, like all database searching methods, is dependent on the reproducibility of the enzyme digestion and both the quality and the completeness of database being searched. The example PCDL database reported here is provided as a demonstration. Due to glycan complexity and the likely absence of specific glycans within the Spike batches prepared by us, it will always be incomplete. Moreover, individual glycopeptides were identified with variable degrees of certainty, and we recommend that they should be validated by the user. As with all glycan analysis methods, there is a bias towards glycopeptides that are easiest to identify by the techniques used, and such bias will also be reflected within the database. Once the PDCL has been created, it must be refined and extended over time to improve data quality, and it is our intention to do so.

## Acknowledgements

The authors wish to thank Professor David Harvey for critical reading of the manuscript and for helpful comments. We thank Professor Ray Owens for kindly providing the RBD-6H construct and Professor Gavin Screaton and Dr. Juthathip Mongkolsapaya for kindly providing the Spike construct. AdvancedBio Peptide HPLC column was a gift from Agilent Technologies. This project has received funding from the Innovative Medicines Initiative 2 Joint Undertaking (JU) under grant agreement No 875510. The JU receives support from the European Union’s Horizon 2020 research and innovation programme and EFPIA and Ontario Institute for Cancer Research, Royal Institution for the Advancement of Learning McGill University, Kungliga Tekniska Hoegskolan, Diamond Light Source Limited. The SGC is a registered charity (number 1097737) that receives funds from AbbVie, Bayer Pharma AG, Boehringer Ingelheim, Canada Foundation for Innovation, Eshelman Institute for Innovation, Genentech, Janssen, Merck KGaA, Darmstadt, Germany, MSD, Ontario Ministry of Research, Innovation and Science (MRIS), Pfizer, São Paulo Research Foundation-FAPESP, Takeda, and Wellcome [106169/ZZ14/Z].

